# A Porcine Model of Peripheral Nerve Injury Enabling Ultra-Long Regenerative Distances: Surgical Approach, Recovery Kinetics, and Clinical Relevance

**DOI:** 10.1101/610147

**Authors:** Justin C. Burrell, Kevin D. Browne, John L. Dutton, Suradip Das, Daniel P. Brown, Franco A. Laimo, Sanford Roberts, Dmitriy Petrov, Zarina Ali, Harry C. Ledebur, Joseph M. Rosen, Hilton M. Kaplan, John A. Wolf, Douglas H. Smith, H. Isaac Chen, D. Kacy Cullen

## Abstract

Approximately 20 million Americans currently experience residual deficits from traumatic peripheral nerve injury. Despite recent advancements in surgical technique, peripheral nerve repair typically results in poor functional outcomes due to prolonged periods of denervation resulting from long regenerative distances coupled with relatively slow rates of axonal regeneration. Development of novel surgical solutions requires valid preclinical models that adequately replicate the key challenges of clinical peripheral nerve injury. Our team has developed a porcine model using Yucatan minipigs that provides an opportunity to investigate peripheral nerve regeneration using different nerves tailored for a specific mechanism of interest, such as (1) nerve modality: motor, sensory, and mixed-modality; (2) injury length: short versus long gap; and (3) total regenerative distance: proximal versus distal injury. Here, we describe a comprehensive porcine model of two challenging clinically relevant procedures for repair of long segmental lesions (≥ 5 cm) – the deep peroneal nerve repaired using a sural nerve autograft and the common peroneal nerve repaired using a saphenous nerve autograft – each featuring ultra-long total regenerative distances (up to 20 cm and 27 cm, respectively) to reach distal targets. This paper includes a detailed characterization of the relevant anatomy, surgical approach/technique, functional/electrophysiological outcomes, and nerve morphometry for baseline and autograft repaired nerves. These porcine models of major peripheral nerve injury are suitable as preclinical, translatable models for evaluating the efficacy, safety, and tolerability of next-generation artificial nerve grafts prior to clinical deployment.

## Introduction

It is estimated that nearly 20 million patients in the U.S alone suffer from chronic deficits as a result of peripheral nerve injury (PNI).^1,2^ Global costs for nerve repair and regeneration are estimated to increase from $5.13 B in 2016 to 10.59 B in 2022.^3^ Nerve injuries, including root avulsions, present in 2–5% of all trauma cases due to various causes, including vehicle accidents, sports-related injury, assaults, combat situations, or iatrogenic incidents.^4–6^ Crush or stretch nerve injuries that do not result in damage to overall nerve structure generally result in a wait-and-see approach to determine if function returns spontaneously.^7–9^ Alternatively, more severe PNI resulting in a disconnection require a surgical procedure to reconnect the proximal and distal nerve stumps by direct anastomosis, a biological or synthetic graft, or nerve conduit.^6^ However, outcomes of surgical repair for traumatic PNI are generally unsatisfactory, irrespective of repair strategy or injury location.^10^

Poor functional outcomes generally stem from long regenerative distances coupled with relatively slow rates of axonal regeneration (~1–2 mm/day), creating prolonged periods of denervation that negatively impact the capacity for axon regeneration as well as the ability of distal nerve structures and their targets to support regeneration and reinnervation, respectively.^10^ Indeed, following severe nerve injury the axon segment distal to the injury site rapidly undergoes Wallerian degeneration with concomitant breakdown of myelin. Regenerating axons from proximal of the injury site must then cross any gap between the proximal and distal nerve stumps, as well as the entire distal nerve segment to reinnervate distal end targets. To facilitate this process, Schwann cells in the proximal and distal nerve undergo choreographed alterations in structure and function, resulting in a transient pro-regenerative phenotype and the formation of regenerative micro-columns called the bands of Büngner. However, the pro-regenerative environment degrades over several months, which may ultimately blunt axonal regeneration while rendering distal sensorimotor targets and/or muscles irrevocably unresponsive to reinnervation, thereby resulting in poor functional recovery.^6,10,11^

Current surgical repair approaches are unable to overcome these challenges. In cases where tension-free direct anastomosis is not possible, the current “gold standard” to bridge a segmental defect remains an autograft repair, which involves deliberately excising an otherwise uninjured nerve of less functional significance (e.g., removing a purely sensory nerve to repair a defect in a motor nerve). The donor nerve acts as a bridge between the proximal and distal nerve stumps by providing a living, Schwann cell laden and matrix-rich scaffold for axonal regeneration. However, long segmental defects (e.g., exceeding the critical gap length of 4–5 cm) and/or proximal (e.g., peri-midline) nerve injuries result in limited sensorimotor functional recovery. Indeed, these two crucial parameters – each presenting a unique set of challenges – must be considered for nerve regeneration: the graft length as well as the total regenerative distance required for functional recovery, as axons must regrow from the transection site, across the graft region, and then within the distal nerve segment(s) to distal targets. Indeed, proximal PNI generally results in diminished functional recovery due to the greater total regenerative distance required to reach distal end targets. Accordingly, there is an urgent need to understand the mechanisms that hinder regeneration and functional recovery when long sections of nerve and/or proximal nerves are repaired.

The development of novel approaches to improve peripheral nerve regeneration requires the use of valid preclinical models that adequately replicate these key challenges of clinical PNI. Large animal models are uniquely able to replicate the large segmental defects and long total regenerative distances necessary to capture the critical biological processes that hinder nerve regeneration in humans. Although previous studies have developed small animal models with a long gap nerve defect, these do not adequately replicate the neurobiological processes in humans and other large mammals. Indeed, the “critical nerve gap” is approximately 4–5 cm in humans as compared to 1.5–3 cm in small animals.^11,13^ Moreover, large animal models can uniquely replicate other critical features such as nerve composition, nerve diameter, and fascicular number/density. To date, large animal PNI models have included the ulnar nerve in primates^14,15^ and inbred pigs,^16^ the median nerve in sheep,^17^ and the peroneal nerve in canines.^18^

Here, we describe the use of Yucatan minipigs to model two challenging clinically relevant nerve injury scenarios – the deep peroneal nerve repaired using a sural nerve autograft and the common peroneal nerve repaired using a saphenous nerve autograft – each featuring long segmental defects (≥ 5 cm) and ultra-long regenerative distances for re-growing axons to reach distal targets (up to 20 cm and 27 cm, respectively). We include a detailed characterization of the relevant anatomy, surgical approach/technique, functional/electrophysiological outcomes, and nerve morphometry. These novel PNI models in minipigs are suitable as preclinical, translatable models that represent the major challenges for repair and assessment of functional recovery experienced in the clinical setting. Nerve repair strategies, including next-generation advanced regenerative therapies such as anisotropic biomaterials, gene therapy, cell-laden scaffolds, and/or tissue engineered constructs, should undergo thorough efficacy, safety, and tolerability testing in an appropriate large animal model prior to clinical deployment to confirm scale-up of the relevant mechanism(s) of action and overall regenerative efficacy, thus ensuring the most promising new strategies are brought to the forefront of clinical treatment.

## Methods

All procedures were approved by the University of Pennsylvania’s Institutional Animal Care and Use Committee and adhered to the guidelines set forth in the NIH Public Health Service Policy on Humane Care and Use of Laboratory Animals (2015).

### Surgical Preparation

This model utilizes young adult Yucatan minipigs 5–7 months of age weighing 30– 40 kg (Sinclair BioResources, Columbia, MO). The Yucatan strain of minipigs was specifically chosen to replicate certain critical features of extremely challenging clinically-relevant repair and regeneration scenarios following major PNI. A total of 14 pigs were specifically utilized to acquire data presented in this study to (a) examine the lower leg nerve anatomy and branching to determine maximum suitable defect lengths (n=3); (b) perform nerve conduction and muscle electrophysiology in naive animals (n=3); (c) evaluate recovery kinetics following repair of a 5 cm deep peroneal nerve lesion using the sural nerve as a sensory nerve autograft (n=4); and (d) evaluate recovery kinetics following repair of a 4 or 5 cm common peroneal nerve lesion using the saphenous nerve as a sensory nerve autograft (n=4).

All surgical procedures were performed under general anesthesia. Animals were anesthetized with an intramuscular injection of ketamine (20–30 mg/kg) and midazolam (0.4–0.6 mg/kg) and maintained on 2.0–2.5% inhaled isoflurane/oxygen at 2 L/min. Preoperative glycopyrrolate (0.01–0.02 mg/kg) was administered subcutaneously to control respiratory secretions. All animals were intubated and positioned in lateral recumbency. An intramuscular injection of meloxicam (0.4 mg/kg) was delivered into the dorsolateral aspect of the gluteal muscle and bupivacaine (1.0–2.0 mg/kg) was administered subcutaneously along the incision site(s) for intra- and post-operative pain management, respectively. The surgical site was prepared and draped under sterile conditions. Heart and respiratory rates, end tidal CO_2_, and temperature were continuously monitored. In our experience, a team consisting of a surgeon, scrub nurse, veterinary anesthetist, and electrophysiologist was adequate for proper execution of these surgeries.

### Anatomical Dissection

Detailed anatomic dissections were carried out in Yucatan cadavers to identify peripheral nerve origins, trajectories, and branching in the hind limbs of this strain of minipigs. A linear incision was made extending from the hip joint to the lateral malleolus. The gluteal muscles were exposed and the inferior gluteal muscle was split longitudinally. The sciatic nerve and its bifurcation into the tibial and common peroneal nerves were identified. The common peroneal nerve (CPN), also known as the common fibular nerve, formed from the lateral aspect of the sciatic nerve, was isolated as it entered the popliteal fossa around the fibular neck until its bifurcation into the superficial peroneal nerve (SPN) and deep peroneal nerve (DPN), also referred to as the superficial and deep fibular nerves, respectively (Figure 1, **middle**). The fascial planes between the extensor digitorum longus (EDL) and the peroneus tertius, and the extensor fibularis longus and fibularis tertius, were dissected to further expose the DPN and its branches (Figure 1, **right**).

**Figure 1.**
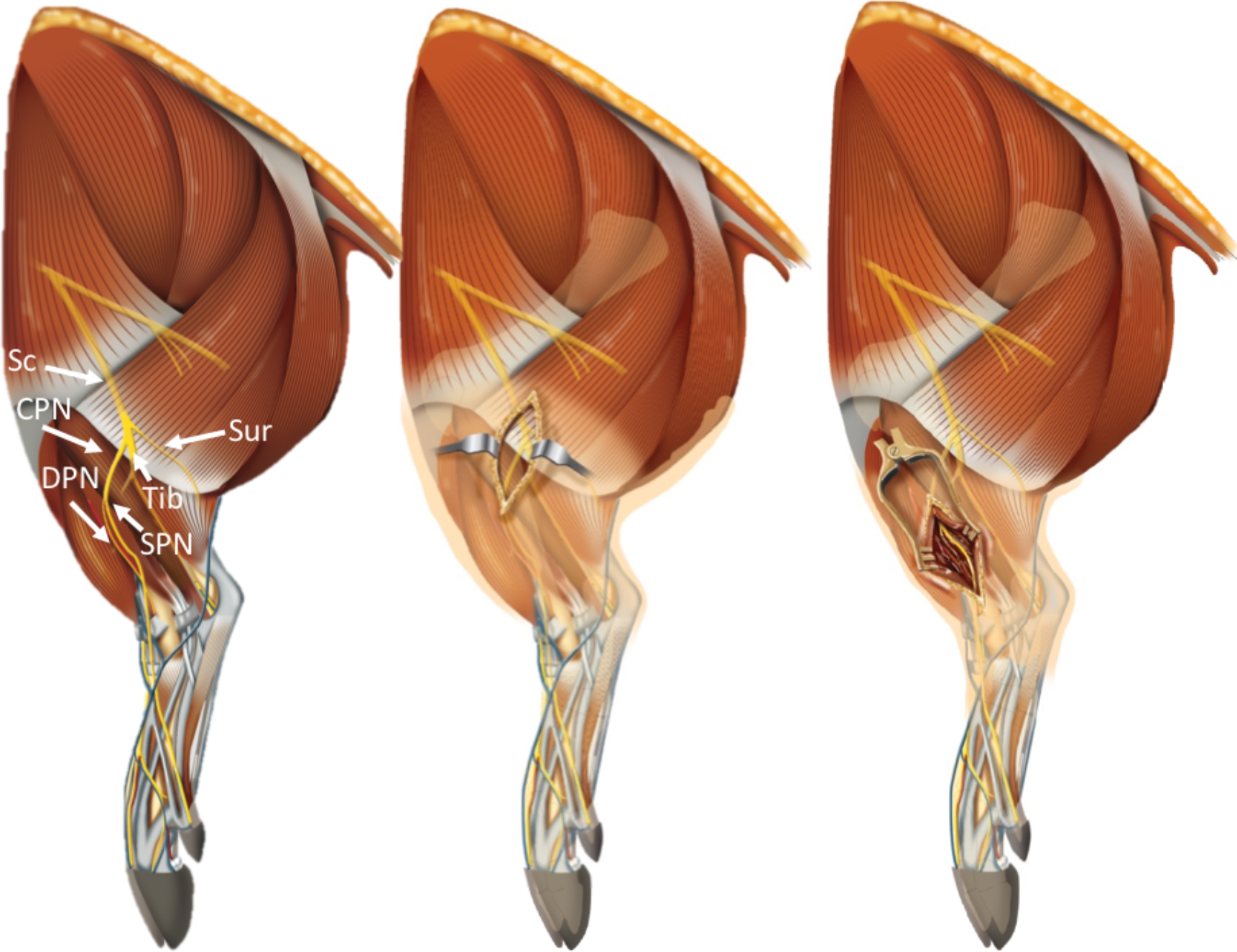
Illustrative View of the Porcine Anatomy and Surgical Approaches to Major Hind Limb Nerves. LEFT: General overview of the A) sciatic nerve (Sc), B) common peroneal nerve (CPN), C) deep peroneal nerve (DPN), D) superficial peroneal nerve (SPN), and E) sural nerve (Su). CENTER: Operative window exposing the junction of the CPN, DPN, and SPN. RIGHT: Operative window exposing the isolated DPN and SPN. Sc and CPN: mixed nerve; DPN: motor nerve; SPN, Su: sensory nerve.

### Operative Technique

The surgical approaches to expose the CPN (mixed motor-sensory) and DPN (mixed motor-sensory), and saphenous (predominantly sensory) and sural nerves (predominantly sensory) for autograft repair of the CPN and DPN, respectively, in Yucatan minipigs are described below.

#### Long Segmental Defect of DPN Repaired with a Sural Nerve Autograft

A 10 cm longitudinal incision was made on the lateral aspect of the right hind limb 1.5 cm distal to the stifle joint and extending to the lateral malleolus (Figure 2A). The fascial layer was bluntly dissected and the peroneus longus was retracted to expose the distal aspect of the CPN diving into the muscle plane between the EDL and tibialis anterior. Further dissection revealed the bifurcation of the DPN and SPN from the CPN (Figure 2B). Distal to the bifurcation, three major branches of the DPN were visualized: a motor branch immediately innervating the tibialis anterior (Figure 2B: **DPN Branch A**), a cutaneous branch coursing superolateral (sensory DPN; Figure 2B: **DPN Branch B**) and motor branch (motor DPN; Figure 2B: **DPN Branch C**) coursing inferomedial to the EDL. Nerve modality was confirmed during exploratory dissections with direct electrical nerve stimulation and visualization of motor movement or lack thereof. Mobilization of the EDL enabled exposure of the entire length of the DPN inferomedial to the EDL extending deep to the flexor retinaculum to innervate the extensor digitorum brevis (EDB). Saline-soaked gauze was placed in the surgical field to prevent desiccation throughout the autograft donor nerve harvest.

**Figure 2.**
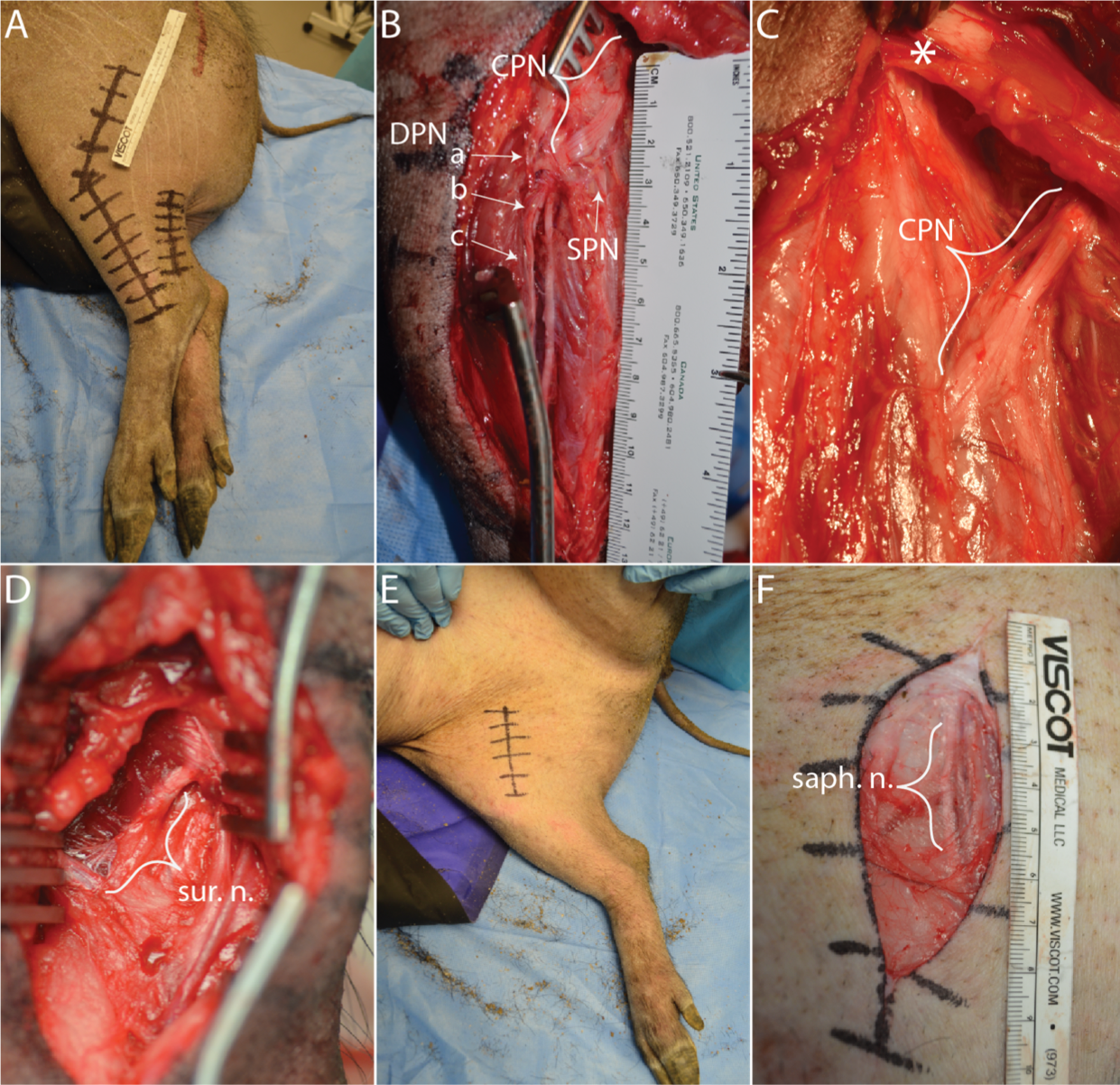
Surgical Approaches to Porcine Hind Limbs. A curvilinear incision was made (A) and the muscle plane was bluntly dissected to reveal the branches of the common peroneal nerve (B). To expose more the proximal aspect of the common peroneal nerve (C), the biceps tendon was cut as denoted with the asterisk. The sural nerve was exposed for the deep peroneal autograft repair (D). The saphenous nerve was exposed through an incision in the inguinal region for the common peroneal autograft repair (E-F). In Panel B, the branches of the DPN are denoted by the innervation point: (a) tibialis anterior motor branch, (b) sensory DPN with cutaneous innervation (b), motor DPN innervating the extensor digitorum brevis (c). **Abbreviations:** DPN–Deep Peroneal Nerve; Sensory DPN–Sensory Branch of the DPN; Motor DPN–Motor Branch of the DPN.

The use of the sural nerve as an autologous donor for repair of the DPN is advantageous from a surgical perspective as there is no need to re-position the animal when accessing the nerve. For harvesting the sural nerve, a second longitudinal incision was made approximately 3 cm posterior to the lateral malleolus and parallel to the Achilles tendon (Figure 2A). The fascial tissue was dissected to expose the sural nerve running close to the saphenous vein, and a 5.5–6.0 cm segment was excised and carefully placed on gauze soaked with sterile saline with its orientation marked to facilitate the reverse autograft repair (Figure 2D). The deep layers and skin were closed with 3-0 vicryl and 2-0 PDS, respectively.

The DPN was carefully dissected from its surrounding tissue and a 5 cm segment was excised, approximately 0.5 cm distal to the bifurcation of the CPN. The harvested sural nerve was trimmed to 5 cm and placed in the surgical field and arranged 180° relative to its normal proximal-distal orientation, for a tensionless reverse autograft with two 8-0 prolene simple interrupted epineural sutures at each end.^19^ The autograft was reversed (i.e. the proximal DPN stump sutured to the distal end of the sural nerve segment, and the distal DPN stump sutured to proximal end of the sural nerve segment) to prevent regenerating axons from entering the distal sural nerve branches.^19^

#### Long Segmental Defect of CPN Repaired with a Saphenous Nerve Autograft

A 4 cm incision in the inguinal region was made to expose the saphenous nerve (Figure 2E). After dissecting the subcutaneous and fascial layer, the gracilis and sartorious muscles were separated to expose a portion of the saphenous nerve (Figure 2F). This exposure enabled an adequately large segment of the nerve to be excised (> 6 cm), which was then carefully placed on gauze soaked with sterile saline with its orientation marked to facilitate the reverse autograft repair.

To expose a suitable length of the CPN, the tendon of the biceps femoris was partially cut to increase exposure of the CPN proximally from approximately 3 cm to 6 cm overall (Figure 2C). Further proximal exposure was limited due to the nerve coursing deeply and closer to its bifurcation from the sciatic nerve. A 4 or 5 cm nerve defect was created and repaired with the saphenous nerve autograft using two 8-0 prolene simple interrupted epineural sutures at each end, resulting in a tensionless repair. Although nerve repairs can be up to 5 cm long within this anatomical location, a tension-less repair was achieved more consistently in grafts of 4 cm in length.

At the conclusion of the harvests/repairs, the areas were irrigated with sterile saline and the fascia and subcutaneous tissues were closed in layers with 3-0 vicryl interrupted sutures. In the CPN cases, the biceps femoris tendon was repaired using 3-0 prolene square suture. The skin was closed with 2-0 PDS interrupted, buried sutures and the area was cleaned and dressed with triple antibiotic ointment, a wound bandage, and a transparent waterproof, adhesive wound dressing.

### Clinical Observations: Motor Deficit

Clinical observations of hoof drop, toe placement, and leg extension were made following surgical repair immediately postoperatively and intermittently thereafter in long-term survival experiments.

### Selection of Terminal Time-Points & Terminal Gross Pathology

Based on our overall experience performing surgical repairs of 5 cm lesions of the DPN (> 40 minipigs) and CPN (> 10 minipigs), the earliest signs of distal muscle reinnervation can be determined at approximately 6- and 9-months post-repair, respectively (unpublished data). Therefore, in order to ensure a more mature and consistent level of functional recovery, the terminal time points that we routinely employ are 9-months for the DPN repair and 12-months for the CPN repair.

### Muscle Electrophysiological Evaluation

Non-invasive compound muscle action potential (CMAP) was recorded from the EDB muscle during transcutaneous nerve stimulation (pulse width: 2.0 ms, amplitude: 0– 10 mA, frequency: 1 Hz) using bipolar bar surface electrodes (2 mm inter-electrode distance; Rochester Electro-Medical, Lutz, FL). A bipolar subdermal recording electrode Medtronic, Jacksonville, FL; #8227410) was placed in the EDB muscle belly, parallel to the muscle fibers, approximately 5 cm distal to the tendon notch on the surface of the hoof. A ground electrode (Medtronic, Jacksonville, FL; #8227103) was inserted into the plantar aspect of the interdigital cleft.

CMAPs were recorded from the EDB following cathodic stimulation (AM Systems, Carlsborg, WA) at two locations two locations: (1) proximal (S1) and (2) distal (S2) to the DPN repair (Figure 3). Proximal stimulation was achieved by approximating the location of the common peroneal nerve coursing near the stifle joint. Distal stimulation was achieved at the hock joint. Once the nerve was located, the stimulus intensity was increased to obtain a supramaximal CMAP (Note: to avoid nerve damage, the absolute maximum stimulus applied was 10 mA at 1 Hz and 0.2 ms). All CMAP recordings were filters. A train of 5 pulses was averaged to increase the signal to noise. The CMAP area under the curve (AUC), the baseline-to-peak amplitude, duration, and latency were measured.

**Figure 3.**
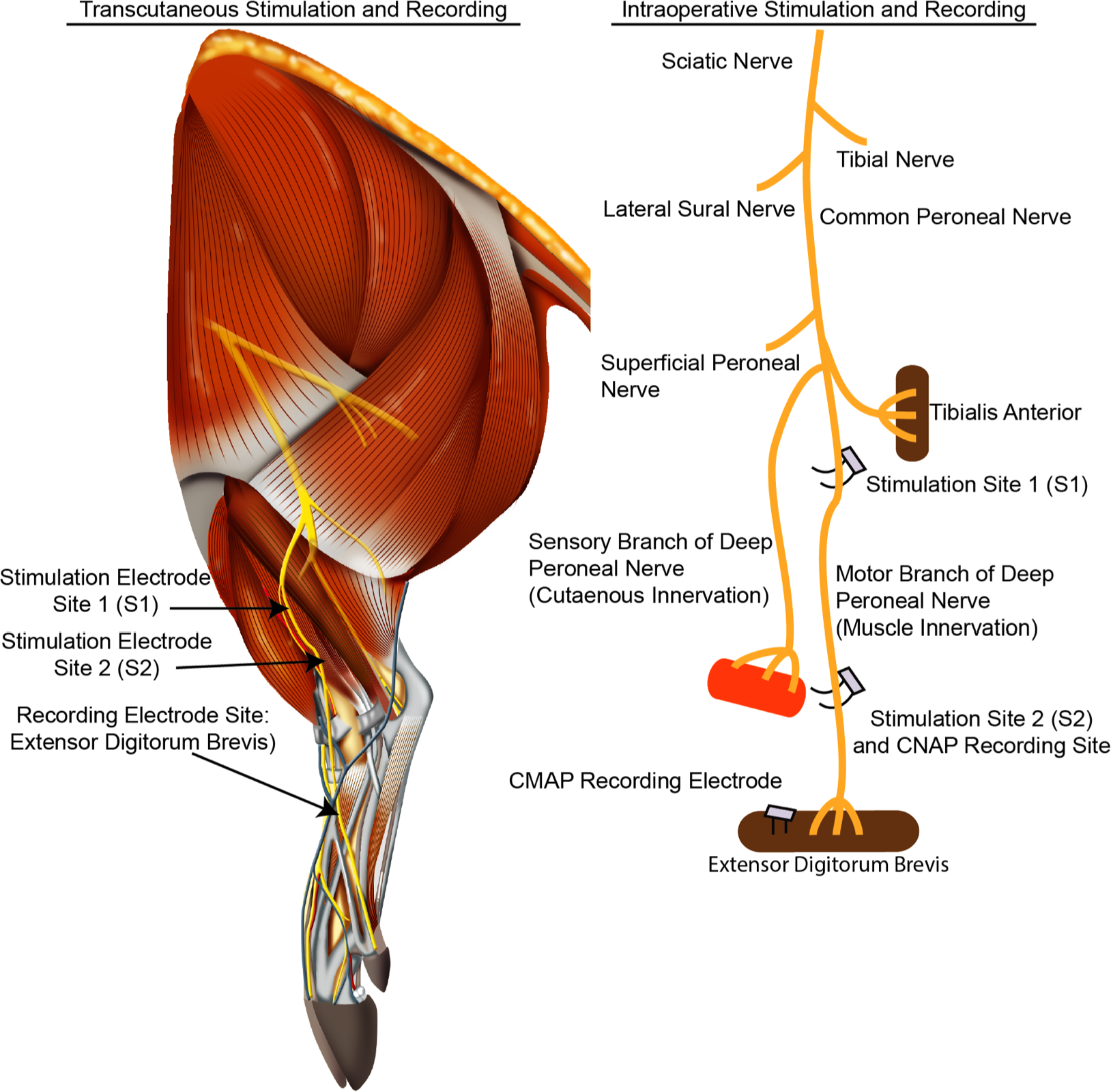
Schematic of the Functional Assessment of the Deep Peroneal Nerve. Electrodes were placed 0.5 cm distal to the bifurcation and 5.5 cm from the bifurcation. Compound nerve action potentials (CNAPs) were recorded by stimulating the electrode proximal to the graft and recording from the electrode distal to the graft. Compound muscle action potentials (CMAPs) were recorded by stimulating proximal and distal to the graft region and recording from the extensor digitorum brevis with subdermal electrodes.

### Nerve Electrophysiological Evaluation

At 9-months post-repair, the DPN was re-exposed and the segment containing the repair was carefully freed to minimize tension and to ensure electrical isolation from surrounding tissue using a rubber mat. The nerve was stimulated 5 mm proximal to the repair zone with a handheld bipolar hook electrode (Rochester Electro-Medical, Lutz, FL; #400900). Compound nerve action potentials (CNAPs) were recorded 5 mm distal to the repair zone with a bipolar electrode with a bend fashioned to maintain better contact with the nerve (Medtronic, Jacksonville, FL; #8227410 – biphasic; amplitude: 0–1 mA; pulse width: 0.2 ms; frequency: 1 Hz; gain: 1000x, bandpass filter: 10–2000 Hz). The ground electrode (Medtronic, Jacksonville, FL; #8227103) was inserted into subcutaneous tissue halfway between the electrodes (Figure 3). A train of 5 pulses were averaged to increase the signal to noise. Mean CNAP peak-to-peak amplitude and latency were calculated.

Baseline CMAP and CNAP measurements were obtained and summarized as naïve data, and contralateral recordings from each animal as internal controls and to calculate percent recovery.

### Euthanasia, Tissue Processing, Histology, and Microscopy

At the conclusion of the functional measurements, all animals were deeply anesthetized (5% isoflurane, 2.5 L oxygen) and transcardially perfused with 4 L heparinized saline followed by 7 L 10% neutral-buffered formalin using a peristaltic pump (ThermoScientific, model #72-320-000). Hind limbs were removed and post-fixed in 10% neutral-buffered formalin at 4 °C overnight to minimize handling artifact resulting under-fixation. The next day, the ipsilateral and contralateral CPN and DPN were isolated and further post-fixed in 10% neutral-buffered formalin at 4 °C overnight.

For histological processing, a 1.0 cm segment of the DPN or CPN, 0.5 cm distal to the repair site, was embedded in paraffin. The block was then mounted on a microtome and sectioned axially at a thickness of 8 µm, mounted on glass slides, and stained for to label neurofilament (a cytoskeletal protein expressed in axon) and myelin (an insulating sheath surrounding mature axons) as follows. Sections were deparaffinized in xylene and rehydrated with a descending gradient of ethanol. Following rehydration, antigen retrieval was performed in TRIS/EDTA buffer for 8 minutes using a modified pressure cooker/microwave technique. Next, normal horse serum in Optimax (Biogenex) was applied to the sections (VectaStain Universal kit per manufacturer’s instructions). Sections were incubated overnight at 4 °C with mouse anti-SMI31/32 (to label neurofilament proteins; Millipore NE1022/NE1023; 1:1000) and chicken anti-myelin basic protein (to label myelin; Encor, CPCA-MBP; 1:1500) in Optimax + normal horse serum (VectaStain Universal kit per manufacturer’s instructions). After washing the sections three times for 5 minutes with PBS/TWEEN, anti-mouse AlexaFluor-567 was applied for 1 hour at room temperature. After rinsing three times for 5 minutes with PBS/TWEEN, was applied for 20 minutes. Finally, sections were washed as above and cover slipped.

Images were obtained with a Nikon A1R confocal microscope (1024×1024 pixels) with a 10x air objective and 60x oil objective using Nikon NIS-Elements AR 3.1.0 (Nikon Instruments, Tokyo, Japan).

## Results

### Characterization of Structure & Function in Naïve Nerves

#### Anatomy and Surgical Approach

The bifurcation of the SPN and DPN from the CPN was located in the superolateral region of the leg between the fibula and peroneus longus. Immediately distal to the bifurcation, a small branch innervating the cranial tibialis was visualized (analogous to the anterior tibialis in humans), and the two major branches of the DPN were identified: (1) a sensory cutaneous branch coursing parallel and superficial to the EDL and (2) a predominantly motor branch of the DPN coursing inferomedial to the EDL extending deep to the flexor retinaculum and innervating the EDB. The motor branch of the DPN (motor DPN) continued deeper into the compartment, parallel to the anterior tibial artery, until coursing under the flexor retinaculum innervating the EDB. A slight ‘s’-shaped bending of the DPN was observed before the nerve entered the anterior compartment inferior to the EDL. The cutaneous branch of the DPN (sensory DPN) descended along the peroneus longus and peroneus tertius, continuing along the fibula before coursing superficially to terminate in the anterolateral aspect of the distal leg near the lateral malleolus.

#### Histological Assessment: Naïve Nerve Morphometry

Normal porcine nerves consisted of large and small myelinated fibers organized in a polyfascicular pattern. Representative images of naïve branches of the DPN and the sural nerve, and the CPN and the saphenous nerve are shown in Figures 4 and 5, respectively. The CPN was the largest of the nerves in this study, and contained a relatively dense fascicular structure. The two major branches of the DPN were similar in diameter and appeared to have a varied density in fascicular structure relative to the total nerve diameter (Figure 4, **top**). The number of fascicles in the DPN (~4–8) closely matched the sural nerve (~4–7) (Figure 4) and the number of fascicles in the CPN (~12–18) closely matched the saphenous nerve (~12–16) (Figure 5).

**Figure 4.**
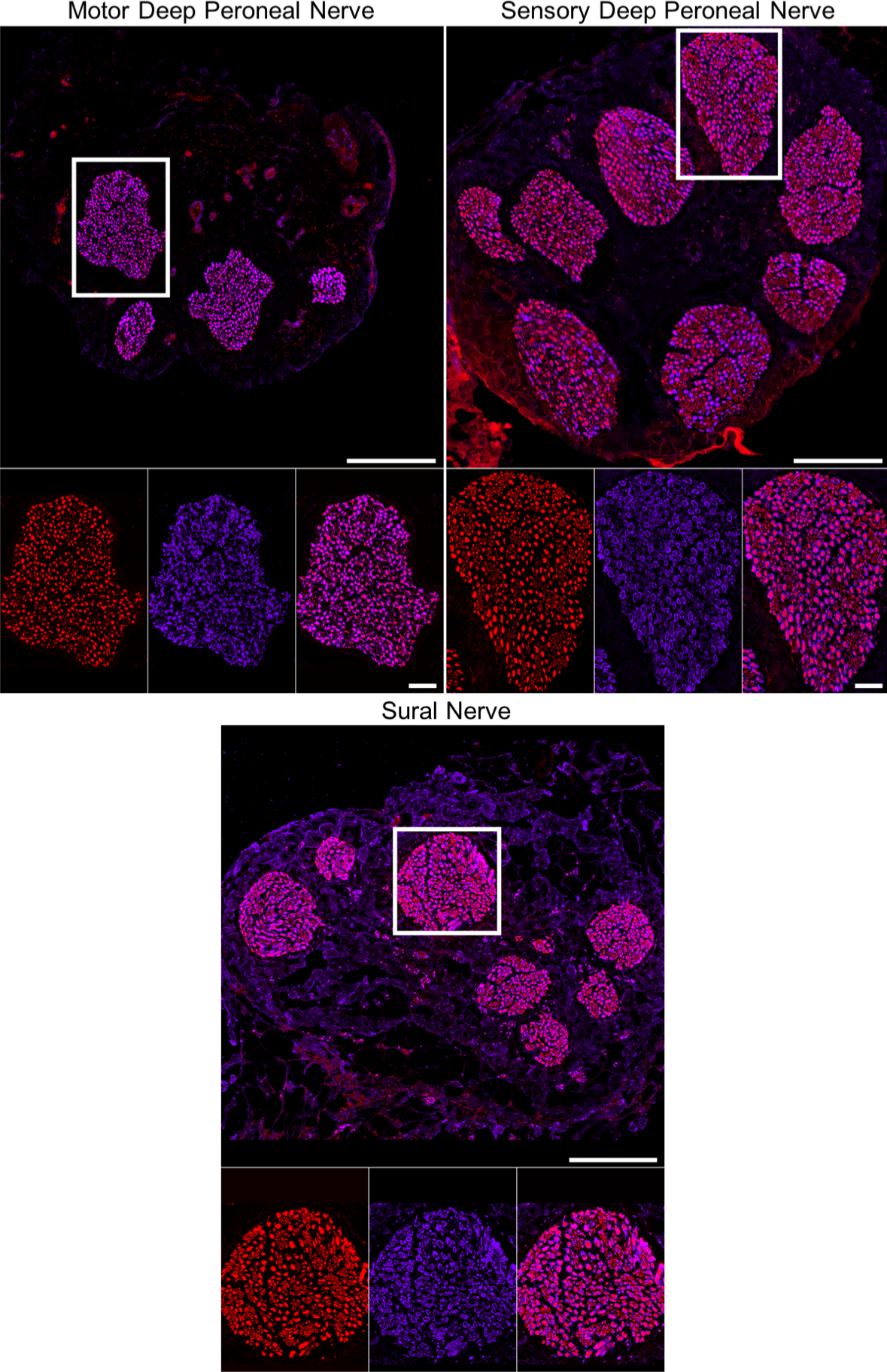
Comparison of Naïve Nerve Architecture of the Motor and Sensory Deep Peroneal Nerves and the Sural Nerve. Naïve motor branch of the DPN (top left), sensory branch of the DPN (top right), and sural nerve (bottom) were stained to label neurofilament (red) to visualize myelinated and unmyelinated axons and myelin basic protein to visualize myelin (purple). Similar polyfascicular organization was found in all nerves. Scale bars - macro: 250 μm; zoom in: 25 μm.

**Figure 5.**
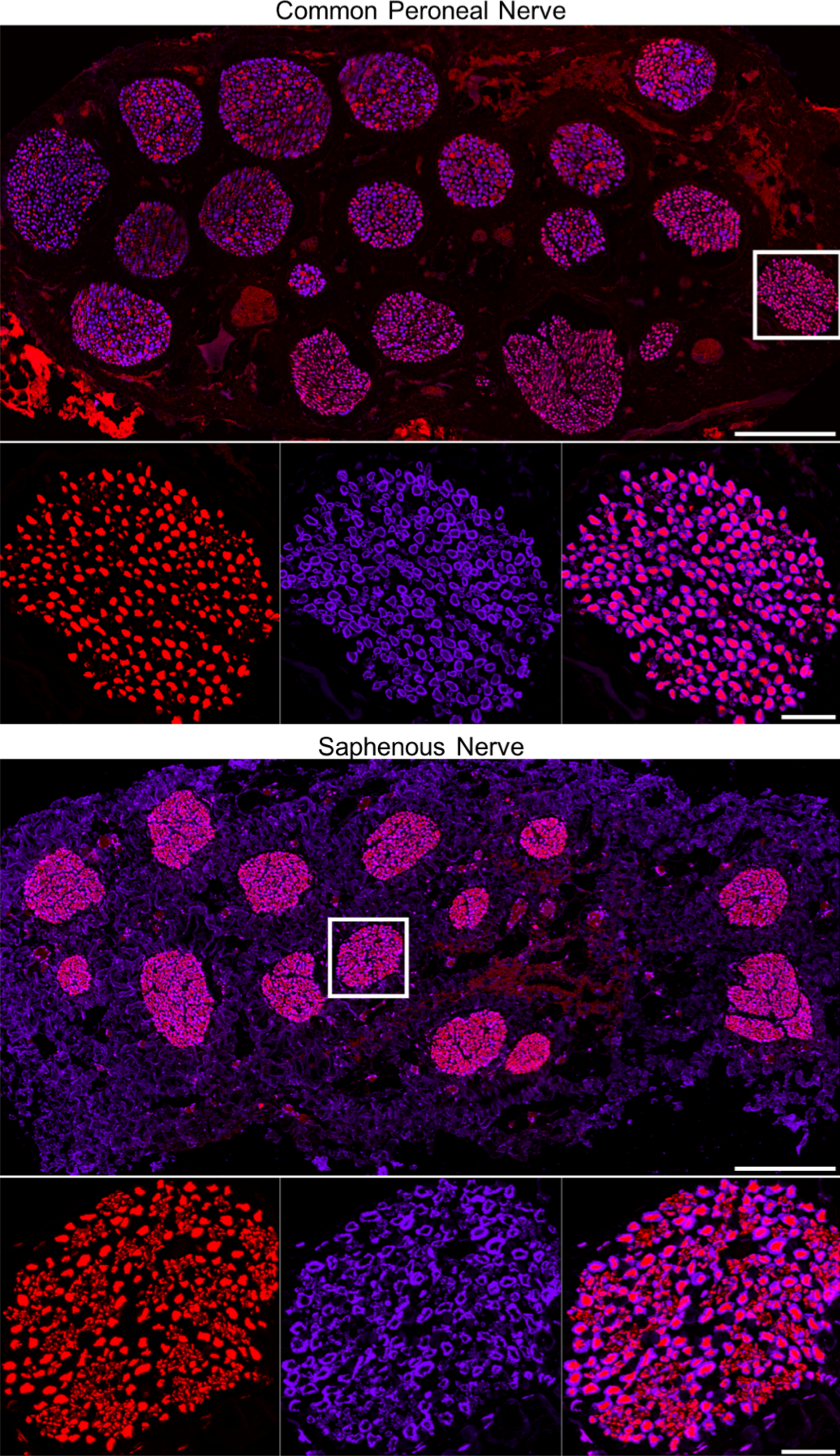
Comparison of Naïve Nerve Architecture of the Common Peroneal Nerve and the Saphenous Nerve. Naïve CPN (top), saphenous nerve (bottom) were stained with neurofilament (red) to visualize myelinated and unmyelinated axons and myelin basic protein to label myelin (purple). Similar polyfascicular organization was found in all nerves. Scale bars - macro image: 250 μm; zoom in: 25 μm.

#### Electrophysiological Functional Assessment: Naïve Nerve and Muscle Recordings

In this study, CMAP peak-to-baseline amplitude, AUC, latency, and duration were measured from the EDB, the distal muscle innervated by the DPN. Proximal nerve and distal nerve electrode placement were standardized for each stimulation paradigm, i.e. surface (transcutaneous) and direct (intraoperative) stimulation. Proximal nerve stimulation resulted in a longer CMAP latency than distal nerve stimulation within each stimulation paradigm, as expected based on the distance from the stimulating electrode. Compared to direct nerve stimulation, indirect activation of the nerve with transcutaneous surface stimulation requires diffuse current to pass through muscle and connective tissue, likely decreasing the effective current that reaches the nerve as well as increasing the latency between the stimulation artifact and CMAP recording. The CMAP duration within each stimulation paradigm was similar for either proximal and distal stimulation.

Mean CMAP and CNAP recordings are summarized with representative waveform traces (Figure 6). Naïve electrophysiological recordings across stimulation methods are summarized in Table 1.

**Table 1.** Normative Deep Peroneal Nerve Electrophysiological Data

| AVERAGE ± SD | Stim. Site | Peak-to-Baseline Amp (mV) | AUC | Lat (ms) | Dur (ms) |
| --- | --- | --- | --- | --- | --- |
| Surface CMAP | S1 | 0.8 ± 0.4 | 1.3 ± 0.5 | 2.6 ± 1.3 | 7.4 ± 1.2 |
|  | S2 | 0.7 ± 0.2 | 1.6 ± 0.6 | 1.5 ± 1.0 | 5.9 ± 1.3 |
| Direct CMAP | S1 | 1.1 ± 0.9 | 2.0 ± 1.9 | 1.8 ± 0.5 | 8.1 ± 0.8 |
|  | S2 | 1.5 ± 1.1 | 3.2 ± 2.5 | 0.7 ± 0.2 | 8.6 ± 1.1 |
|  |  | Peak-to-Peak Amp (mV) | Lat (ms) |  |  |
| CNAP |  | 2.7 ± 1.3 | 1.3 ± 0.4 |  |  |

**Abbreviations:**
SD: Standard Deviation; Stim: Stimulation; AUC: Area-Under-Curve; Lat: Latency, Dur: Duration; CMAP: Compound Muscle Action Potential; CNAP: Compound Nerve Action Potential.

**Figure 6.**
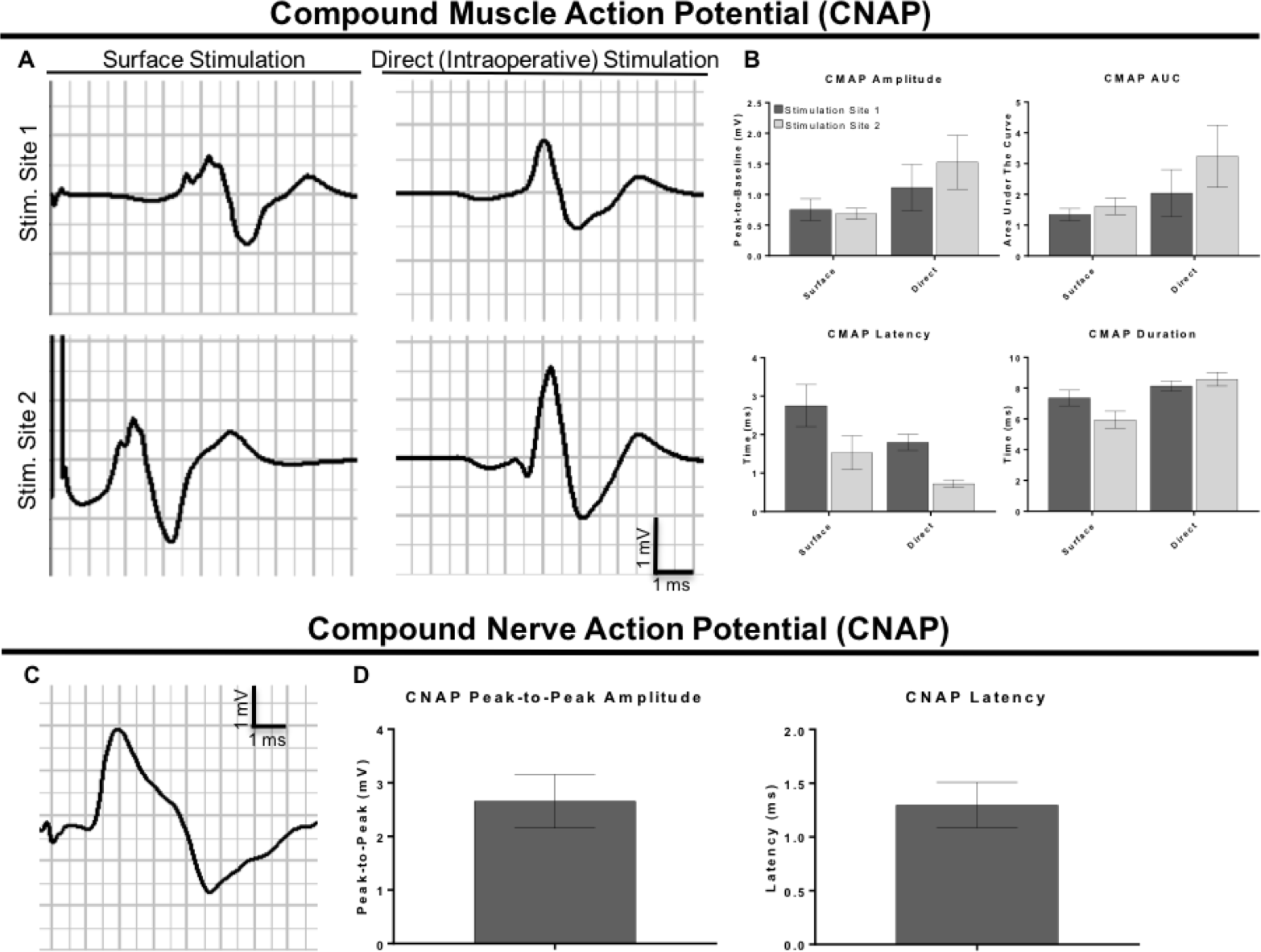
Comparison of Various Electrophysiological Functional Assessment Methods. Compound muscle action potential (CMAP) recordings from the EDB were evoked with surface (transcutaneous), percutaneous or intraoperative nerve stimulation (A). Mean CMAP amplitude, area-under-the-curve (AUC), latency, and duration are shown for each CMAP stimulation paradigm (B). Compound nerve action potential (CNAP) recordings were obtained by stimulating the nerve and recording the electrical activity in the distal region (A). Peak-to-peak amplitude and latency were measured (B). Representative waveforms were averaged over a train of 5 pulses are shown with the starting time denoting the end of the stimulus artifact.

### Functional & Structural Metrics of Regeneration Following DPN Surgical Repair

#### Surgical Trauma & Repair of the DPN

In the surgical approach described above, the branches of the DPN were visualized with the sensory branch coursing approximately 8 cm from its origin towards the lateral malleolus, typically the location near the cutaneous innervation site; and the motor branch extending in a deeper plane to course under the flexor retinaculum, approximately 7–8 cm from its origin. For terminal investigations, the flexor retinaculum can be sharply divided to increase the distal limit of the surgical window and provide greater exposure for electrophysiological experiments. The total regenerative distance across the long gap nerve injury and to the distal muscle end target of the motor DPN in this study was approximately ~20 cm. For 5 cm autograft repairs, 6 cm of the sural nerve was excised; however, longer lengths are available if needed. The diameter of the sural nerve was comparable to the diameter of the DPN (Figure 7; also shown in Figure 4).

**Figure 7.**
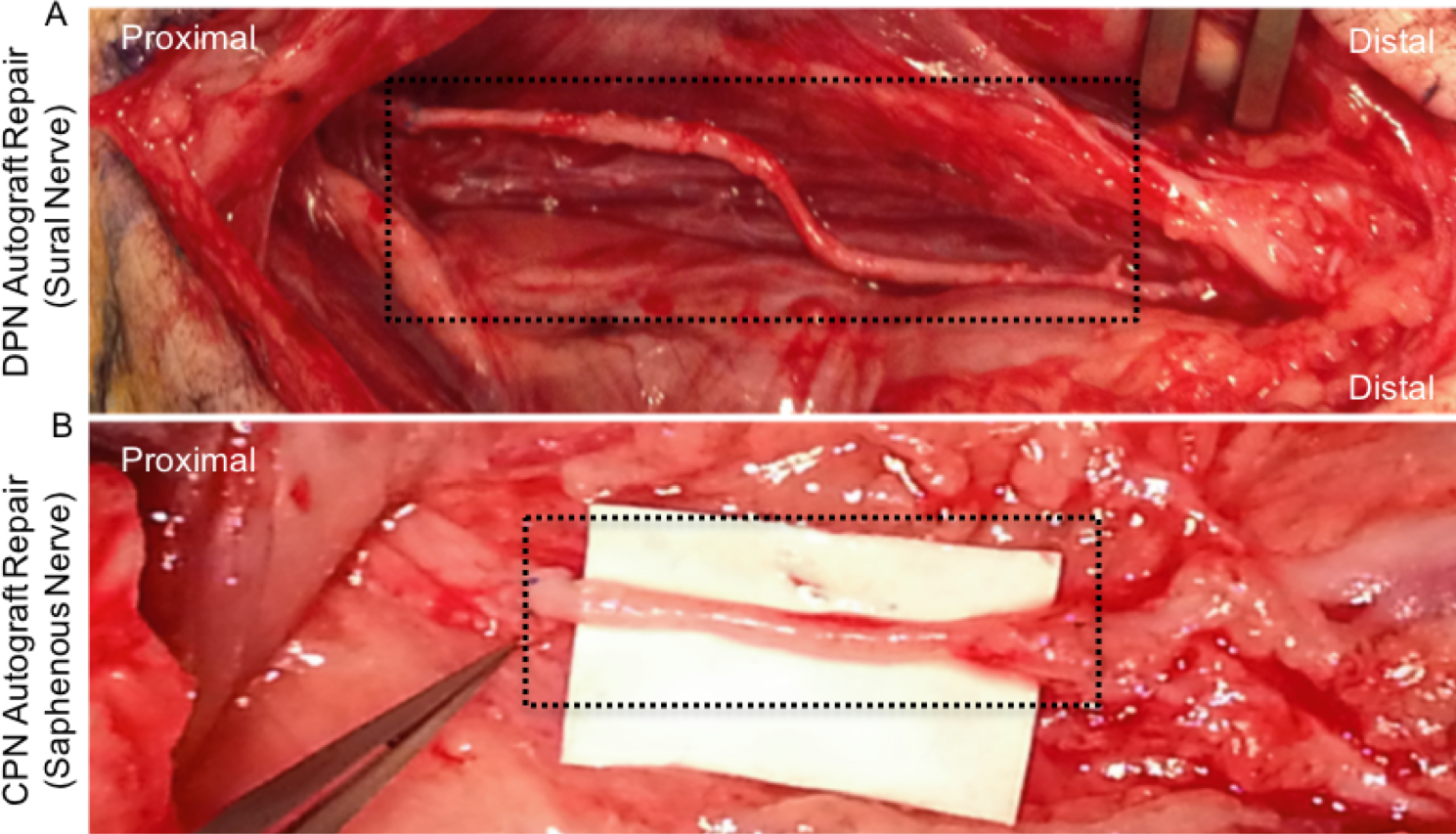
Representative Images of Long-Gap Nerve Injury Repaired Nerves. (A) In this example, the deep peroneal nerve was repaired with a 5 cm sural nerve autograft (Top black dotted box). (B) In this example, the common peroneal nerve was repaired with a 4 cm saphenous nerve autograft (Bottom black dotted box).

#### Functional Deficits of the DPN

Transection of the individual nerves resulted in different clinical outcomes. Transection of the motor DPN resulted in noticeable hoof drop whereas transection of the sensory DPN had no directly observable motor deficits. All animals with a transected DPN had normal limb extension and ambulation with full weight-bearing capacity within 1–2 hours of recovery from anesthesia. As expected, the animals favored the use of the contralateral limb after surgery; however, this resolved within 24 hours. After a few weeks, minimal changes in gait were observed including decreased ipsilateral toe placement and minor hoof drop. We did not observe further favoritism or compensatory actions during the experimental period and all animals were fully weight bearing and ambulatory. Although lameness in the affected limb became less apparent over time, muscle atrophy was appreciable at 6- and 9-months post-repair. There were no surgical complications such as infection, suture dehiscence, or prolonged swelling as a result of the surgical procedure.

#### Terminal Gross Pathology of the DPN

The graft region was visualized by locating the permanent prolene sutures demarcating the proximal and distal coaptation sites. At the terminal time point, minimal fibrosis surrounding the autograft repair was observed. Healthy vascularization of the regenerated nerve was achieved in all repaired nerves. The nerve tapered slightly at the interface between the host nerve stumps and the graft location, likely because of the small diameter mismatch between the DPN and sural nerve autograft, and the degree of regeneration.

#### Electrophysiological Functional Recovery at 9-Months Post-Repair of the DPN

To evaluate functional recovery of the DPN at 9-months post-repair using an autograft, CMAP recordings were obtained from the EDB. The surface stimulation and direct nerve stimulation paradigms were used to evaluate whether non-invasive stimulation and intraoperative stimulation resulted in similar electrophysiological outcomes. Representative CMAP and CNAP waveforms and mean peak-to-baseline amplitude, latency, AUC, and duration values, and percent recovery are shown in Figure 8.

**Figure 8.**
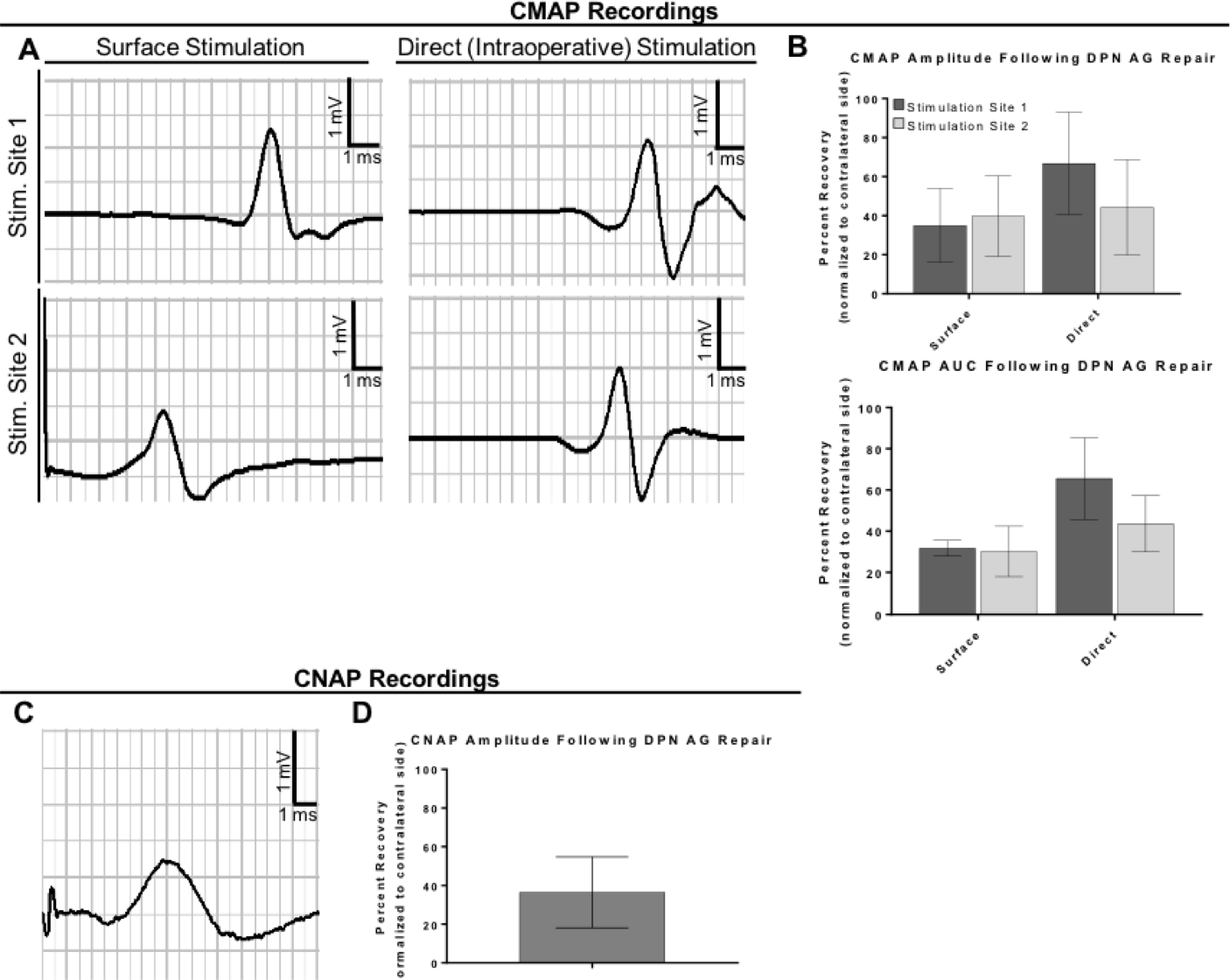
Electrophysiological Functional Recovery at 9-Months Post-Repair. All repaired nerves elicited robust CMAPs at 9-months post-repair (A). Mean CMAP Amplitude and AUC are shown for the surface and intraoperative nerve stimulation paradigms (B). An injury effect was demonstrated by calculating the percent recovered by normalizing the values measured for the repaired nerve (ipsilateral) to the naïve side (contralateral) for each animal. All repaired nerves elicited robust CNAPs at 9 months post-repair (C). Mean peak-to-peak Amplitude and Percent Recovery are presented (D). An injury effect was demonstrated by calculating the percent recovered by normalizing the values measured for the repaired nerve (ipsilateral) to the naïve side (contralateral) for each animal.

The injury effect was readily measured with both stimulation paradigms. However, the degree of recovery appeared to be greater with direct nerve stimulation than surface stimulation. It is likely that this is due to direct activation of more regenerating axons and corresponding increased recruitment of muscle fibers. CNAP recordings measured the electrical conduction of axons that had regenerated across the graft zone. Restoration of electrical conduction corroborated the muscle electrophysiological data.

#### Nerve Morphometry at 9-Months Post-Repair

In concordance with the electrophysiological functional recovery, regenerated axons were found distal to the repair site at 9-months post-repair (Figure 9). Large and small caliber axons in the distal nerve were organized in a polyfascicular structure similar to the naïve architecture at 9-months post-repair (Figure 9). Myelination around a portion of the regenerated axons indicated ongoing maturation, and small myelinated fibers were also visualized across the various fascicles. These finding demonstrate that numerous axons successfully crossed the 5 cm sural nerve autograft, with at least a portion of them ultimately traveling a total regenerative distance of ~20 cm to reinnervate the EDB.

**Figure 9.**
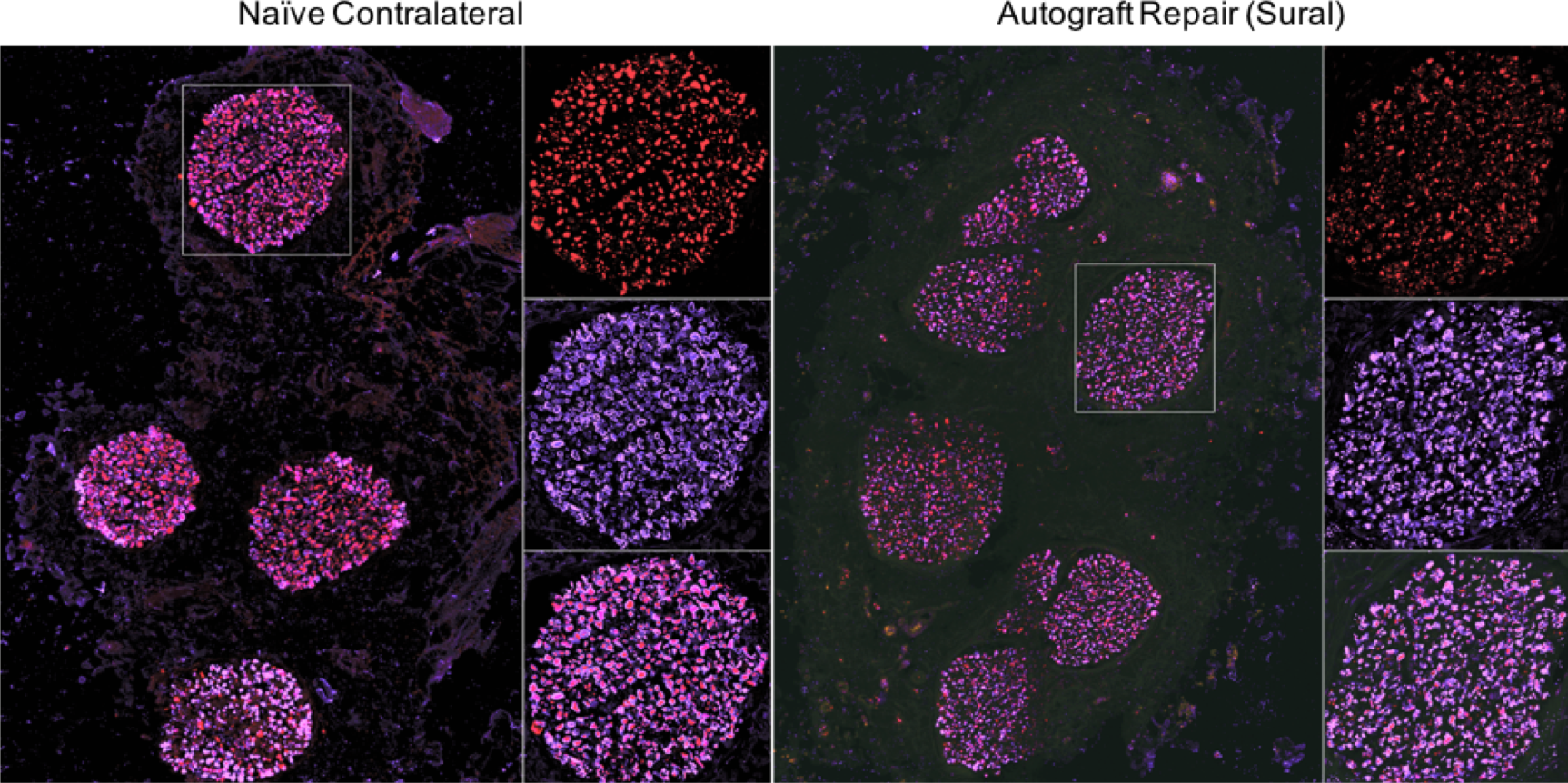
Comparison of Naïve (Contralateral) Motor DPN and Motor DPN Repaired Using a Sural Nerve Autograft at 9-Months Post-Repair. Confocal reconstruction of the deep peroneal nerve, 5 mm distal to the 5 cm repair site. Regenerated nerves (right) as compared to naïve contralateral nerves (left) were labeled to denote neurofilament (red) and myelin (purple). Scale bars - macro image: 250 μm; zoom: 25 μm.

### Functional & Structural Metrics of Regeneration Following CPN Surgical Repair

#### Surgical Trauma & Repair of the CPN

In the surgical approach described above, approximately 3 cm of the CPN was exposed. Greater exposure could be achieved, up to approximately 6 cm by dividing the tendon of the biceps femoris and further retraction. For 4–5 cm autograft repairs, 6 cm of the saphenous nerves could be excised; however, longer lengths are available if needed. The diameter of the saphenous nerve was a close match to the diameter of the CPN (see Figure 7; also shown in Figure 5). Surgical transection and repair of the CPN in this manner results in ~25 cm total regenerative distance being required to reinnervate distal muscle end-targets.

#### Functional Deficits of the CPN

The surgical approach to expose the CPN resulted in inability of full hind limb extension during the acute post-operative phase, likely associated with tendon dissection. The animals were able to stand with full weight-bearing capacity typically within 3–4 hours. As expected, the animals favored the use of the contralateral limb after surgery; however, this resolved within 24 hours. After a few weeks, minimal changes in gait were observed including decreased ipsilateral toe placement and minor hoof drop. We did not observe further favoritism or compensatory actions during the experimental period and all animals were fully weight bearing and ambulatory. Lameness in the affected limb became less apparent over time. Muscle atrophy of the EDB and tibialis anterior was apparent at 6-, 9-, and 12-months post-repair. There were no surgical complications such as infection, suture dehiscence, or prolonged swelling as a result of the surgical procedure.

#### Terminal Gross Pathology of the CPN

The graft region was visualized by locating the permanent prolene sutures demarcating proximal and distal coaptation sites. At the terminal time point, minimal fibrosis surrounding the autograft repair was observed. Healthy vascularization of the regenerated nerve was achieved in all repaired nerves. The nerve generally slightly tapered at interface between the host nerve stumps and the graft location, likely because of the small diameter mismatch between the CPN and saphenous nerve autograft.

#### Electrophysiologic Functional Recovery at 9- and 12-Months Post-Repair of the CPN

To evaluate the extent of nerve regeneration following CPN autograft repair, functional recovery was assessed at 9- and 12-months post-repair. Representative CMAP and CNAP waveforms are shown in Figure 10. A small CMAP was observable at 9-months post-repair, indicating regenerating axons had crossed the graft region begun to reinnervate the distal EDB muscle, approximately 27 cm from the injury. Robust electrophysiological recordings were obtained at 12-months post-repair, indicating further nerve regeneration than at 9-months post-repair likely due to maturation and myelination of the motor axons innervating the distal muscle target. Indeed, CNAP recordings were measured from the two branches distal to the CPN defect, indicating nerve regeneration resulted in the restoration of electrical conduction.

**Figure 10.**
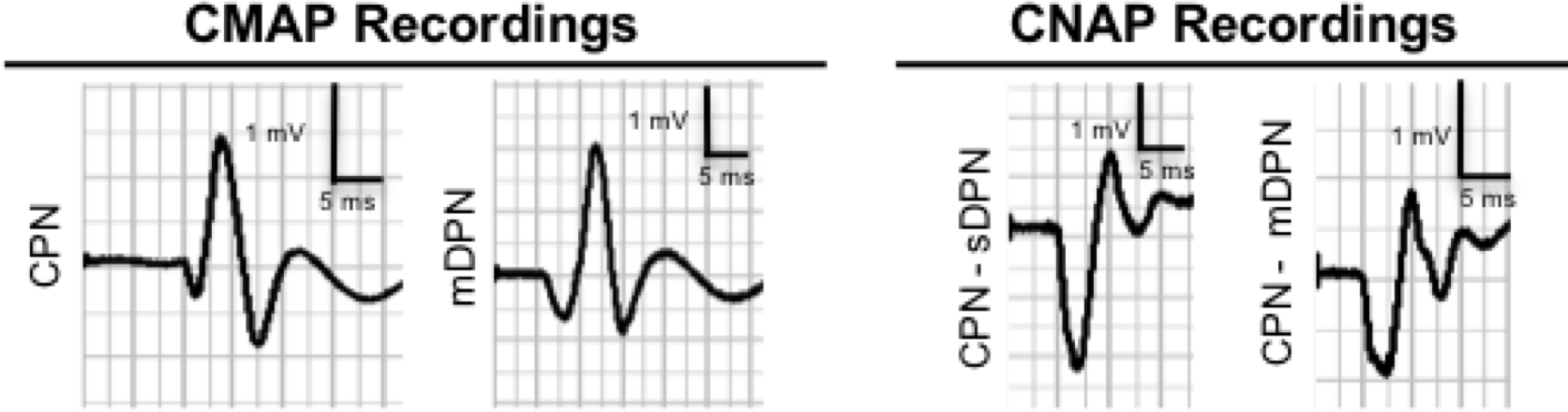
Example Electrophysiological Functional Recovery Following CPN Repair Using a Saphenous Nerve Autograft at 12 Months Post-Repair. Representative electrophysiological traces are shown. At 9-months post-repair, CMAP were recorded by stimulating proximal to the nerve defect. (Scale bars. 05 mV / 6 ms). At 12-months post-repair, CMAPs were recorded stimulating proximal to the repair site (CPN) and distal to the repair site from the motor DPN branch. CNAPs were also recorded across the CPN graft from the sensory DPN and motor DPN branches (Scale bars 1 mV / 6 ms).

#### Nerve Morphometry at 12-Months Post-Repair of the CPN

Representative histological data are shown in Figure 11. As with the repaired DPN nerve, a polyfascicular pattern of small and large, myelinated and unmyelinated axons was found in the CPN, distal to the autograft at 12-months post-repair. In this model, axons successfully spanned the 4-5 cm saphenous nerve autograft repair, with evidence of distal penetration of these regenerating axons, in support of the functional findings.

**Figure 11.**
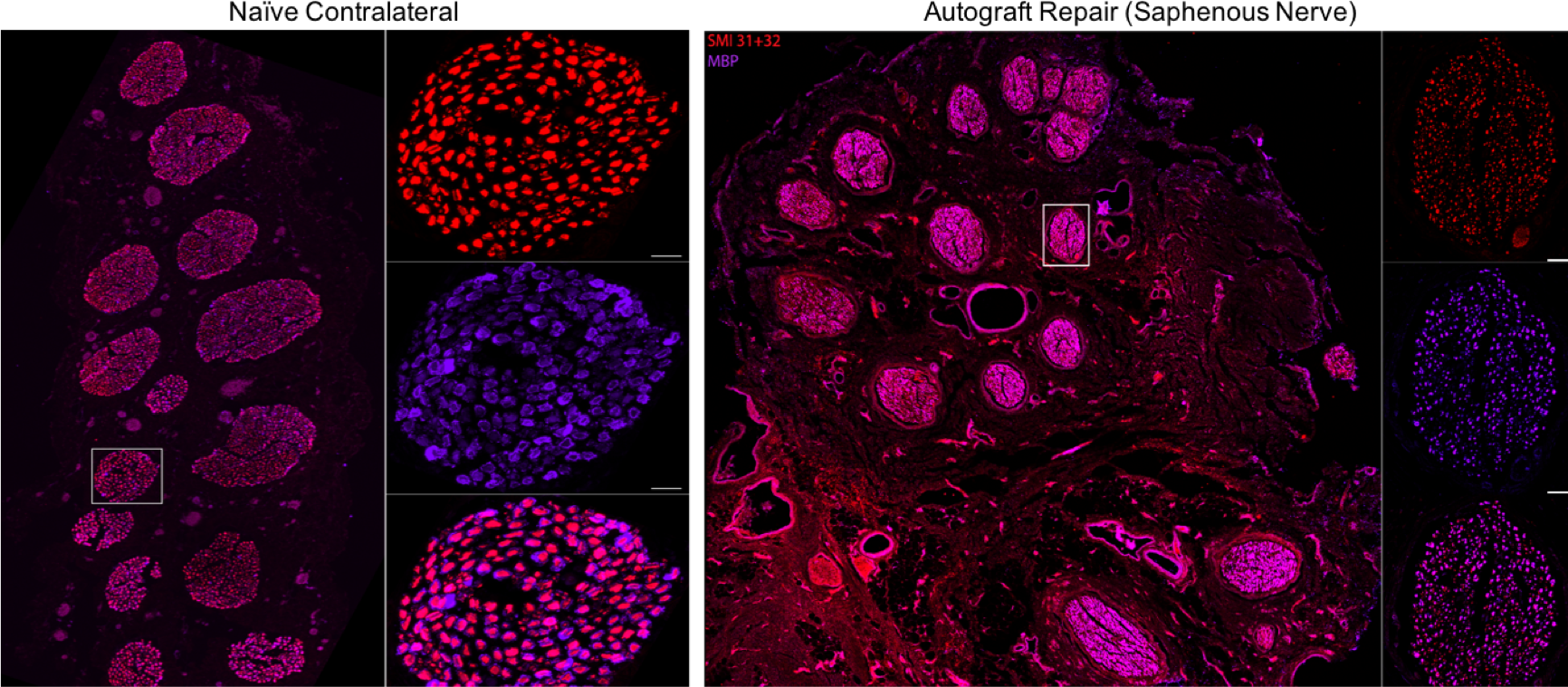
Comparison of Naïve (Contralateral) CPN and CPN Repaired Using a Saphenous Nerve Autograft at 12-Months Post-Repair. Confocal reconstruction of the common peroneal nerve, 5 mm distal to the repair site. Regenerated nerve (right) was compared to naïve contralateral nerves (left) following labeling to denote axons (red) and myelin (purple). Scale bars - macro image: 250 μm; zoom: 25 μm.

## Discussion

In this study, we described the surgical approaches and recovery kinetics for two long-gap nerve injury models that mimics the common clinically relevant scenario of using sensory nerve autografts for repairing motor and/or mixed motor-sensory nerves. Here, we evaluated regeneration across 4-5 cm lesions; however, greater distances are attainable with more muscle dissection (CPN: up to 7 cm, DPN: up to 8–10 cm; Table 2). Alternative complex neurosurgical procedures are possible using this model, such as multiple nerve repairs using a “stepping stone technique”, nerve transfers that enables innervation of an damaged nerve from a neighboring uninjured, ultra-long segmental deficit starting at the proximal CPN to the distal DPN (> 15 cm), or electrical stimulation to enhance regeneration.^20–27^ Previous studies have developed small animal models as an attempt to replicate challenging clinical scenarios, such as long gap nerve injury, in lower order species.^28–31^ However, we believe that these models do not adequately replicate the neurobiological processes found in humans and other large mammals, thus highlighting a major concern in the evaluation of potential clinical products in rodent models.

**Table 2.** Exposed Nerve Lengths, Recommended Repair Lengths, and Maximum Regenerative Distance from Proximal Injury Site to Distal Muscle End-Target

| **Nerve** | **Modality** | **Nerve Length**<br>**(Surgical**<br>**Exposure)** | **Maximum Repair**<br>**Length**<br>**(Recommended)** | **Maximum Regenerative**<br>**Distance** |
| --- | --- | --- | --- | --- |
| **CPN** | Mixed Motor-Sensory | 7 cm | 5 cm | Up to 27 cm |
| **DPN** | Mixed Motor-Sensory | 8–10 cm | 10 cm | Up to 20 cm |
| **Sur. N** | Predominantly Sensory | Up to 8 cm | N/A | N/A |
| **Saph. N.** | Predominantly Sensory | Up to 8 cm | N/A | N/A |

Nerve regeneration in large animal models, such as dogs, cats, sheep, pigs, and monkeys have been evaluated across nerve gaps ranging from 1 to 9 cm.^32^ Choosing the species in which to model long-distance PNI requires careful consideration. The species must be manageable, cost-effective, amenable to behavioral measurements, and expendable for tissue analysis. Unlike other widely used models, porcine subjects are particularly well suited as a pre-clinical model because the nerves are remarkably similar to humans.^33^ Moreover, the similar physiology between humans and porcine have resulted in its increasing popularity as a preclinical translational research model in various areas, such as traumatic brain injury,^34–40^ spinal cord injury,^41^ wound healing,^42^ cartilage repair,^43–47^ intervertebral disc herniation,^48^ coronary artery injury,^49^ and gastrointestinal,^50^ and hepatic surgery.^51^ Indeed, the objective in the development of the porcine model presented in this study was to replicate critical features of extremely challenging repair and regeneration scenarios following major PNI in human.

Although small animal models remain useful in proof-of-concept experiments such as the mechanism of action and first-order optimization of putative regenerative enhancements (e.g., scaffolds, growth factors, etc.), large animal models are necessary for translational research such as chronic efficacy and tolerability studies including reinnervation following ultra-long distance axonal regeneration and biocompatibility/ immunological responses (Figure 12). The degree that mechanisms of peripheral nerve trauma, degeneration, and regeneration are conserved across species remains unclear.^11,32^ While regeneration studies in rodents offer some utility (higher throughput with lower associated costs), standardized models of long nerve gap injury are necessary to assess treatment strategies following major PNI. Large animals are uniquely able to replicate the large segmental defects (≥ 5 cm) and long total regenerative distances (≥ 20 cm) that are the primary challenges to nerve regeneration and functional restoration in humans.^52^

**Figure 12.**
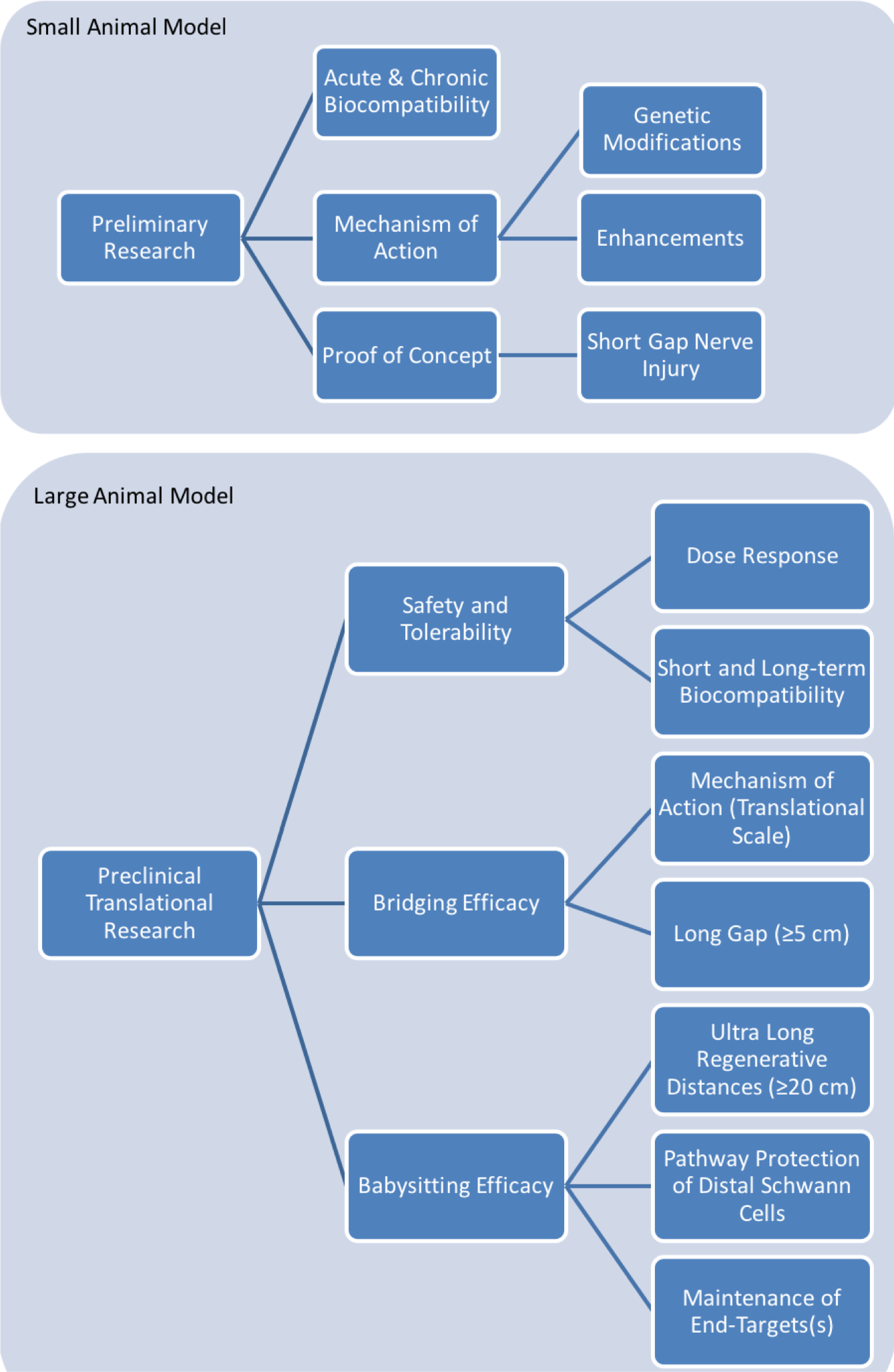
Example Applications for Utilizing Small and Large Animal Model in Nerve Regeneration Research

Within a given species, strain selection is another important consideration to maximize the cost-effectiveness and translatability of the model. Porcine strains currently available for research include Yorkshire and Landrace farm pigs, and several minipig breeds such as Yucatan, Hanford, Sinclair, and Göettigen minipigs.^53^ Young adult Yucatan minipigs weighing approximately 20–30 kg can grow to 50 kg at 9 months of age and will reach their maximum height and weight of ~70–75 kg around 1 year of age. In contrast, Yorkshire pigs will triple or quadruple their size within the first year, reaching almost 150– 200 kg at two years old. On the other end of the spectrum, the average maximum weight for Göettigen minipigs is roughly 40–50 kg.^53^ Our lab has used domestic farm pigs (Yorkshire strain) for short-term studies, e.g. investigating the translatability of various tissue engineered nerve grafts from the rodent to porcine model and the mechanism of action of nerve regeneration following short-gap repair at acute time points. In chronic experiments (> 1 month), such as investigating the efficacy of nerve grafts in long-gap repair, the rate and extent of muscle reinnervation and functional recovery is evaluated in young adult Yucatan minipigs. In this study, the Yucatan minipig strain was selected for our model of nerve regeneration because compared to other farm strains, minipigs have sufficiently long limbs (20–30 cm) and a manageable weight gain to facilitate handling at chronic time points. In addition, ease of access and animal recovery time should be considered when designing a given experiment. In our experience, the nerve and musculoskeletal systems appear relatively similar between Yorkshire and Yucatan strains.

Similar to the human nervous system, the constitution of large nerves of the hind limb is variable (e.g., sensory versus motor constituency, fascicular density, diameter, etc.) and nerves should be chosen based on the experimental question. For example, if a behavioral/motor outcome is desired, it may be advantageous to select a predominantly motor nerve (e.g., motor branch of DPN). Indeed, in this study, the CPN and motor branch of the DPN were chosen as the templates for this comprehensive model of long-gap nerve injury to mimic the clinically relevant scenario using a sensory nerve autograft resulting in motor deficit. On the other hand, if behavioral/motor outcomes are not necessary, alternative nerves, such as the sensory branch of the DPN can also suitable to evaluate general axonal regeneration, maturation, and/or myelination.

Sensory nerve autografts are commonly used in research to compare the effect of various experimental strategies. In this study, our criteria for selecting an autograft donor nerve was a sensory nerve that closely matched the diameter of the injured nerve. In our anatomical study, we found that the diameter of the sural nerve was similar the DPN but about half that of the CPN. On the other hand, the diameter of the saphenous nerve—a large-caliber sensory nerve in the lower leg—was similar to the diameter of the CPN. Moreover, the number of fascicles in the CPN closely matched the saphenous nerve and the number fascicles in the branches of the DPN closely matched the sural nerve. Therefore, the sural nerve was selected as the donor nerve for DPN autograft repair and the saphenous nerve for CPN autograft repair.

Alternatively, surgeons often use nerve cable grafts, or multiple autograft segments used in parallel, to reconstruct large caliber mixed or motor nerves in a clinical situation. In the models presented here, we utilized a single autograft nerve repair to reduce surgical repair variability, reduce donor site morbidity related to having to harvest longer nerve segments, reversed the autograft 180° (i.e. the proximal DPN stump sutured to the distal end of the sural nerve segment, and the distal DPN stump sutured to proximal end of the sural nerve segment) to minimize branching.^19^ Also, in considering nerve and species selection for other applications, human-like fascicular organization is an important consideration for evaluation of peripheral nerve electrical interface devices, suggesting the benefits of a standardized large animal model with polyfascicular nerve architecture in this increasingly common application as well.^55^

Clinically, the CPN is the most commonly injured peripheral nerve in the lower limb,^56^ thus easily providing justification when choosing this nerve for PNI research. While the limbs in most large animals do not have the same complexity as humans, porcine exhibit “hoof drop” comparable to “foot drop” observable in clinical cases of common or deep peroneal nerve. Indeed, transection of the nerves in the hind limb allows residual innervation from spared nerve-muscle groups to maintain weight bearing but with noticeable gait deficits, manifesting as the “hoof drop”. These deficits derived from these hind limb models are amenable to quantitative software-based gait-analyses to assess recovery over time, although these measures are time and resource intensive in this porcine model. Moreover, deficits vary with time and compensatory mechanisms. For instance, animals with common peroneal injuries have limited extension of the limb for a period of time following surgery. Similarly, animals that have undergone deep peroneal nerve injuries will have a noticeable “hoof drop” that can be measured. While these deficits are obvious during acute recovery, as time progresses, the deficits become less noticeable but still present. This may be due to redundancy in innervation or to the possibility that the animals are compensating for their injury. Importantly, muscle atrophy becomes readily apparent and easily palpable. Finally, hind limb repair in a quadruped has the advantage of not affecting a limb supporting the weight of the chest and head, thus making the procedure less traumatic for the animal and limiting aftercare needs and/or other special requirements.

The model presented here provides a framework for evaluating experimental strategies to repair long gap nerve injury with functional regeneration (e.g., 6–12 months). This line of research is not only important for human clinical scenarios, but also for nerve regeneration in veterinary medicine, such as in race horse laryngeal nerve hemiplegia.^57^ Moreover, large animal models are uniquely suitable for the development of new techniques to longitudinally assess nerve injury and regeneration. Previous studies have reported the importance of investigating differences in sensory and motor axon regeneration.^58–64^ Here, we demonstrate that proximal nerve injury to the CPN results in degeneration of a motor and sensory branch of the DPN, enabling researchers to investigate modality-specific axon regeneration in a larger animal model. Indirect measurements to evaluate the severity of nerve injury and monitor the extent of nerve regeneration, such as end-target muscle atrophy, can be performed using ultrasonography or magnetic resonance imaging.^65,66^ Direct measurements of nerve regeneration can be visualized using ultrasonography^67,68^ or diffusion tensor imaging-based tractography.^69–74^ Nerve reinnervation and functional recovery can also be assessed using either direct nerve stimulation by re-exposing the nerve or non-invasively using surface electrodes to stimulate the nerve and record muscle activity. Indeed, previous studies have evaluated regeneration in pigs using ultrasonography, magnetic resonance and diffusion tensor imaging-based tractography, sensory and motor evoked potentials, nerve conduction velocity, muscle activation, and nerve tract tracing.^75–81^

Electrophysiological functional assessments are frequently used to indirectly evaluate nerve regeneration and the extent of target muscle functionality at specific time points.^82–85^ The evaluation and interpretation of functional recovery via nerve electrical stimulation and nerve/muscle recording can be technically challenging due to hardware issues, over- or under-sampling data, stimulus artifact, anatomical variation, or poor functional recovery.^86^ Despite the popularity, comparison of function recovery across studies is often challenging due to a lack of standardized parameters for electrical stimulation (e.g., duration, constant current or voltage, electrode polarity, etc.) and recording parameters/locations.^52,86–89^ Moreover, interpretation of electrophysiological functional recovery is difficult in small animal models: the stimulation of a rodent sciatic nerve can cause inadvertent direct or indirect stimulation of nearby nerves and muscles, which might increase cross-talk at the recording site that resembles physiological CMAPs.^90^ Conversely, large animal models have the advantage of providing a greater distance between stimulation and recording sites, limiting the presence of confounding and/or artifactual signals. Moreover, direct stimulation and recording of nerve activity introduces the possibility of iatrogenic damage to the intact or newly repaired nerve. As an alternative, non-invasive methods can be used such as those described here, designed to match clinical practices as closely as possible. In this model, transcutaneous surface stimulation resulted in muscle activation; however, the muscle and connective tissue surrounding the nerve likely diminished the effective current, and increased the latency between the stimulation pulse and onset of action potentials. It is also possible to avoid complications from multiple invasive procedures by utilizing implantable devices capable of wireless electrical stimulation and recording.^91,92^ We found that these methods were sensitive and reliable in permitting the tracking of axonal regeneration and reinnervation over time without the use of multiple invasive procedures. However, as a final measure, compound muscle and nerve action potentials resulting from intraoperative (direct) nerve stimulation were evaluated at the terminal time point.

In this model, we aimed to develop an intraoperative electrophysiological assessment that minimized the stimulus artifact by maximizing the distance between electrodes and avoiding unnecessary dissection of the nerve. Moreover, a bipolar or tripolar stimulating electrode reduces the onset stimulus artifact that can distort CNAP recordings across short distances (< 4 cm). The location of the stimulating and recording electrodes are standardized relative to the anatomical landmarks, such as the bifurcation of the CPN, and can accommodate grafts up to 5 cm in length. Moreover, in initial evaluations to develop the method for recording CNAPs across the DPN, we found that complete nerve isolation greatly increased the stimulus artifact saturation. In contrast, leaving the nerve buried in connective tissue and limiting the dissection to only the actual stimulating and recording sites decreased stimulus artifact and lessened the likelihood of iatrogenic nerve damage due to tissue dissection. In addition, no previous studies have reported the relationship between the stimulus artifact saturation during CNAP recording and the nerve dissection technique in large animals. In contrast to rodents yet similar to human anatomy, porcine nerves are typically insulated by connective tissue, which might decrease the electrical spread and stimulus artifact during CNAP recordings.

With the advent of neural tissue engineering, many attempts have been made to enhance nerve regeneration, yet the autograft repair still remains the gold-standard of practice.^93^ We assert that next-generation tissue engineered constructs should undergo thorough efficacy, safety, and tolerability testing in large animal models prior to clinical deployment, and that this novel porcine model of major PNI is suitable to represent the major challenges for repair and functional recovery experienced in the clinical setting.

## Conclusions

This article described a reliable and straightforward preclinical model of major PNI and repair in Yucatan minipigs, including nerve selection, surgical approach, motor deficits, functional outcomes, and nerve morphometry. This model may provide a useful platform for scientists and clinicians to evaluate promising next-generation repair strategies. While small animal models are most commonly used in peripheral nerve research due to their ability to achieve higher throughput screening of the regenerative efficacy of various tissue engineered constructs, rodent nerves are smaller in diameter with fascicular organization and extracellular matrix composition unlike analogous human nerves.^33^ Porcine subjects provide an excellent opportunity to investigate peripheral nerve regeneration using different nerves tailored for a specific mechanism of interest, such as (1) nerve modality: motor, sensory, mixed-modality; (2) injury length: short versus long gap; and (3) total regenerative distance: proximal versus distal injury. Unlike other large animals, such as sheep and non-human primates, minipigs are a more cost-effective alternative that requires less space for housing and are easily trained to perform measurable behavioral outcomes. Moreover, minipigs minimize the challenges and confounds associated with the rapid vertical growth often seen in other species. In addition, porcine handling and long-term management may be less cumbersome than that of sheep, canine, or non-human primates. Our study provides detailed information on nerve access, anatomy, composition, and functionality, which we believe will encourage other researchers to consider the use of porcine as clinically-relevant PNI models, especially when bringing new strategies to the forefront of clinical treatment.

## Acknowledgements

The authors acknowledge Laura Struzyna, Kritika Katiyar, Chelsea Wallace, Joseph Maggiore, Gabe DeSantis, and Vishal Tien for technical contributions, and the University Laboratory Animal Resources staff at the University of Pennsylvania for their animal care. Financial support provided by the U.S. Department of Defense [CDMRP/JPC8-CRMRP #W81XWH-16-1-0796 (Cullen), MRMC #W81XWH-15-1-0466 (Cullen), JWMRP #W81XWH-13-207004 (Cullen)], Department of Veterans Affairs [Career Development Awards #IK2-RX001479 (Wolf) and #IK2-RX002013 (Chen)], and American Association of Neurological Surgeons and Congress of Neurological Surgeons [Codman Fellowship in Neurotrauma and Critical Care (Petrov)].

